# Conformational dynamics in TRPV1 channels reported by an encoded coumarin amino acid

**DOI:** 10.1101/137778

**Authors:** Ximena Steinberg, Marina A. Kasimova, Deny Cabezas-Bratesco, Jason Galpin, Ernesto Ladrón-de-Guevara, Federica Villa, Vincenzo Carnevale, Leon D. Islas, Christopher A. Ahern, Sebastian E. Brauchi

**Author notes:** These authors contribute equally (Ximena Steinberg, Marina A. Kasimova). Correspondence should be addressed to Dr. Sebastian Brauchi.

## Abstract

Transient Receptor Potential Vanilloid (TRPV1) channels support the detection and integration of nociceptive input. Currently available functional and structural data suggest that that TRPV1 channels have two potential gates within their cation selective permeation pathway: a barrier formed by a ‘bundle crossing’ at the intracellular entrance and a second constriction created by the ion selectivity filter. To describe conformational changes associated with channel gating within the pore, the fluorescent non-canonical amino acid (f- ncAA) coumarin-tyrosine was genetically encoded at Y671, a residue proximal to the selectivity filter. TRPV1 channels expressing coumarin at either site displayed normal voltage- and agonist-dependent gating. Next, total internal reflection microscopy (TIRF) was performed to enable ultra-rapid, millisecond imaging of the conformational dynamics in single TRPV1 channels in live cells. Here, the data obtained from channels expressed in human derived cells show that optical fluctuations, photon counts, and variance of noise analysis from Y671 coumarin encoded in TRPV1 tetramers correlates closely with channel activation by capsaicin, thus providing an direct optical marker of channel activation at the selectivity filter. In companion molecular dynamics simulations, Y671 displays alternating solvent exposure between the closed and open states, giving support to the optical data. These calculations further suggest a direct involvement of Y671 in controlling the relative position of the pore helix and its role in supporting ionic conductance at the TRPV1 selectivity filter.

## INTRODUCTION

Cation channels from the TRPV family are functional tetramers with a central pore domain that is shaped by TM5 and TM6 transmembrane segments from each of the four subunits (Ramsey et al., 2006) (Figure 1a). As in other *Shaker*-related tetrameric ion channels, the pore domain houses the ion selectivity filter and intracellular bundle crossing formed by the channels S6 segments. Functional and structural data suggest that the selectivity filter and inner bundle crossing have the potential to control the flux of ions according to the electrochemical gradient (Cao et al., 2013; Gao et al., 2016; Salazar et al., 2009). While the lower constriction is highly conserved (Palovcak et al., 2015) and associated to the canonical gate (Steinberg et al., 2014), less is known about the putative role of the upper constriction at the ionic filter, possibly owing the sensitive nature of this region to side-chain mutagenesis. Nevertheless, high resolution structures of TRPV1 suggest that the upper constriction undergoes subtle structural changes in the presence of the DxTx (Gao et al., 2016), a spider toxin that increases the open probability of the channel (Siemens et al., 2006). However, it is not known if such conformational dynamics at the selectivity filter might be coupled to channel activation by agonists, such as capsaicin, and if filter gating is sufficient, or acts in concert with the lower bundle crossing to control ionic conductance.

Organic dyes with environmentally-sensitive fluorescent have been used for nearly 20 years to describe membrane protein conformational changes and ion channel motions (Mannuzzu et al., 1996). The voltage-clamp fluorometry approach (VCF) relies on the covalent attachment of such fluorophores, usually via introduced cysteine residues, and allows for the simultaneous recording of fluorescence signals and voltage-clamped ionic currents from expressed ion channels, often in the Xenopus oocyte (Cha and Bezanilla, 1997). This technique has proven to be useful for the real-time description of subtle, and in some cases electrophysiologically silent, conformational changes in voltage- and ligand-gated ion channels (Cha et al., 1999; Islas and Zagotta, 2006; Pless and Lynch, 2009). However, excessive background labeling of intracellular sites, at endogenous cysteine residues for instance, and the prerequisite for solvent accessibility of the labeling site, serve to substantially limit the utility of the technique for the study of intracellular or solvent restricted residues. A possible solution to these limitations is the use of genetically encoded fluorophores in the form of non-canonical amino acids (f-ncAA) (Drabkin et al., 1996). Such strategies employ an orthogonal suppressor tRNA and an evolved tRNA synthetase (RS), which can be used to encode the ncAA at any site in the reading frame of the target gene (Drabkin et al., 1996; Sakamoto et al., 2002). This approach has been recently used for site-specific incorporation of fluorescent ncAAs on soluble proteins in prokaryotic systems (Summerer et al., 2006; Wang et al., 2006), membrane proteins expressed in Xenopus oocytes (Kalstrup and Blunck, 2013; Pantoja et al., 2009), and proteins expressed in mammalian cells (Chatterjee et al., 2013; Luo et al., 2014; Shen et al., 2011; Steinberg et al., 2016; Zagotta et al., 2016). The f-ncAA coumarin has potential for the study of expressed protein conformational dynamics because of its small size and its exquisite sensitivity to environmental polarity (Liu et al., 2015; Wagner, 2009; Wang et al., 2006). The later can be exploited to report on variations in the surroundings (i.e. dielectric) of the incorporated f-ncAA and, in turn, it can be interpreted as a signature of a protein conformational change. Notable examples include the use of coumarin and the amber codon suppression technique to report on peptide binding to viral proteins (Ugwumba et al., 2011), the photoregulation of firefly luciferase and the subcellular localization of proteins in living mammalian cells (Luo et al., 2014), and structural rearrangements in isolated NaK channels (Liu et al., 2015).

Here, encoded hydroxy-coumarin was used with total internal reflection (TIRF) microscopy and rapid imaging of single emitters to study the conformational changes occurring within the conducting pore of the capsaicin receptor TRPV1 during activation.

## RESULTS

### Expression of Tyr-coumarin at genetically encoded positions

Based on previous cysteine accessibility experiments (Salazar et al., 2009), we reasoned that a genetically encoded coumarin residue at position Y671 (Y671^Coum^) would be functionally tolerated and if so, could potentially would be well positioned to report on local structural rearrangements occurring within the pore during gating (figure 1a). As expected for an encoded fluorescent probe, HEK-293T cells heterologously expressing both TRPV1-YFP and amber codon-containing TRPV1 mutants (TRPV1^YFP^/TRPV1^TAG^) exhibit coumarin-YFP colocalization (figure 1b) consistent with the presence of coumarin and YFP co-fluorescence. Whole cell patch clamp electrophysiological recordings performed at room temperature (22°C) demonstrate the functional expression of the coumarin-containing TRPV1 channels (TRPV1^TAG^) when expressed alone or at 1:7 molar ratio (TRPV1^TAG^:TRPV1-YFP) (figure 1c). The later condition was chosen to simplify the imaging single emitters (i.e. one coumarin molecule per channel tetramer or cluster of tetramers). Macroscopic channel kinetics were similar between WT and coumarin-containing channels (figure 1c) and the conductance-voltage (G-V) relationship from wild type and coumarin channels were similar and well fit by a Boltzmann distribution with a half-maximal activation voltage (V_0.5_) near +100 mV, (figure 1d). Thus, the data suggest that position Y671 is amenable to nonsense suppression and encoding of f-ncAA coumarin.

### Optical recordings of capsaicin-induced changes in solvation

In order to demonstrate the ability of the genetically encoded hydroxy-coumarin side chain to act as reporter for protein activity in live cells, the evanescent field of HEK Y671-coumarin TRPV1 expression cells was imaged with a single-photon avalanche diode (SPAD) photon counting camera (64×32 pixels array with in-pixel counters) (Bronzi et al., 2014; Michalet et al., 2013). Specifically, photon counts of the emission signal were examined to determine the effect of capsaicin, the canonical agonist of the TRPV1 channel (figure 1 e-g). The photon count distribution showed clear differences between pixels representing background and pixels from fluorescent spots, which presumably arise from fluorescent channel subunits (figure 1g). Further, when the cell is exposed to a maximal concentration of capsaicin (3 μM), the photon counting rate further shifted to higher values (figure 1g). One interpretation of the capsaicin-induced increase in photon emission counts is that it stems from local changes in the immediate environment of the dye.

To simplify the analysis and interpretation of the optical data, we set out to directly measure the fluctuations of the fluorescence signal emitted from a single diffraction-limited emitter. Given the large pixel size and the low fill-factor of the SPAD array used (Bronzi et al., 2014; Michalet et al., 2013), the precise localization of a diffraction-limited signal is not possible under the condition of overexpression of membrane proteins in living cells. Therefore, steady state fluctuations of fluorescence were recoded by an EM-CCD camera to resolve changes in the optical properties of the dye (Blunck et al., 2008). As in the electrophysiological experiments, cDNAs encoding for both wild-type channels (TRPV1) and amber codon containing mutants were mixed in 7:1 ratio to promote the formation of uneven channel tetramers. This strategy results in a small yet discrete number of the diffraction-limited coumarin emitting spots as observed by total internal reflection of fluorescence (TIRF) microscopy (figure 2a). The signal colocalize well with the YFP signal corresponding to wild-type YFP-tagged channel subunits (figure 2a). Close examination of these puncta by photobleaching, showed a stepwise behavior allowing for ex-post identification of puncta containing a single emitter (figure 2b). A clear change in fluorescence was evidenced by the ensemble average of several diffraction-limited spots once the cell is exposed to capsaicin (3 μM; figure 2c). The ensemble recording not only showed an increase of fluorescence after agonist treatment, but also increases the standard deviation of the signal’s mean value (figure 2c).

**Figure 1.**
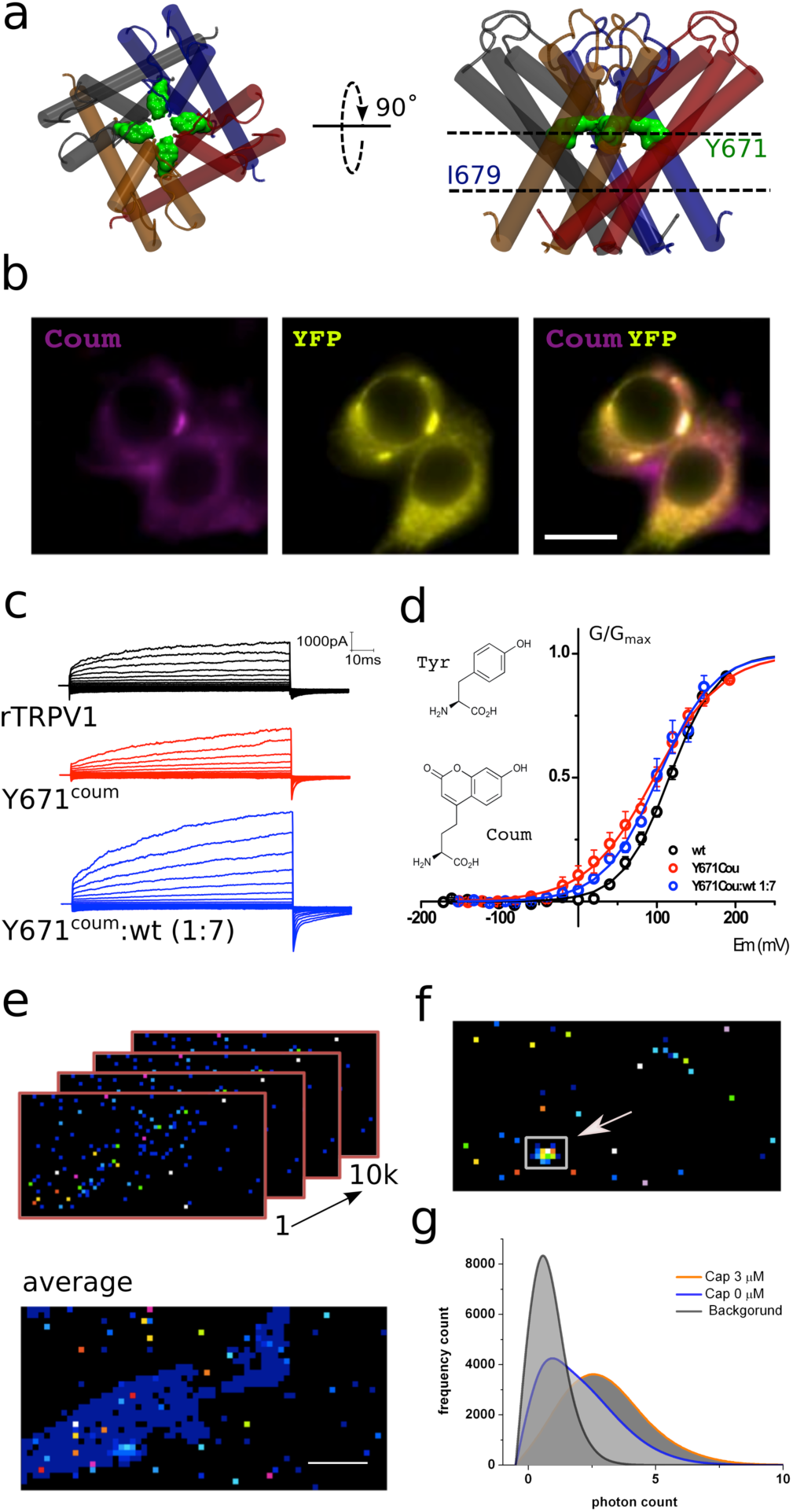
Functional expression of coumarin-containing TRPV1 channels at the membrane of HEK-293T cells. **a.** Top-down and lateral view of the pore region of rTRPV1 tetramers. Dashed lines denote putative gates, a canonical gate at I679 and a secondary constriction within the pore at Y671 (green shade). The different subunits are shown in different colors. **b.** Epifluorescence images obtained from transiently transfected HEK-293T cells. Cells were co-transfected with wild-type rTRPV1-YPF and rTRPV1- Y671^coum^. Bar corresponds to 10 μm. Images correspond to the average of 50 frames. **c.** Representative traces obtained from transiently transfected HEK-293T cells subjected to voltage steps from -180 mV to +180 mV, in 20 mV increments at the three different conditions indicated. **d.** Conductance-voltage relation for the wild-type rTRPV1 (black) and the two different coumarin-containing channels (TRPV1-coumarin alone in red; 1xTRPV1coumarin +7xTRPV1wt in blue). Curves were fitted to a Boltzmann distribution of the form I=I_max_/(1+ exp([zF(V-V_0.5_)/RT]), where z is the voltage dependency, V_0.5_ is the half-activation voltage, and I_max_ is the maximum current. Each curve represents the average of at least four different experiments performed at 20°C. Error bars correspond to SEM. **e.** 10 thousand individual frames were taken with the 64×32 SPAD camera (upper panel) and averaged to identify cell area and ROIs. **f.** A threshold was set on the averaged image in order to define ROIs (arrow and square). **g**. Photon count was extracted from the original set of frames. Capsaicin incubation promotes a higher photon count distribution.

**Figure 2.**
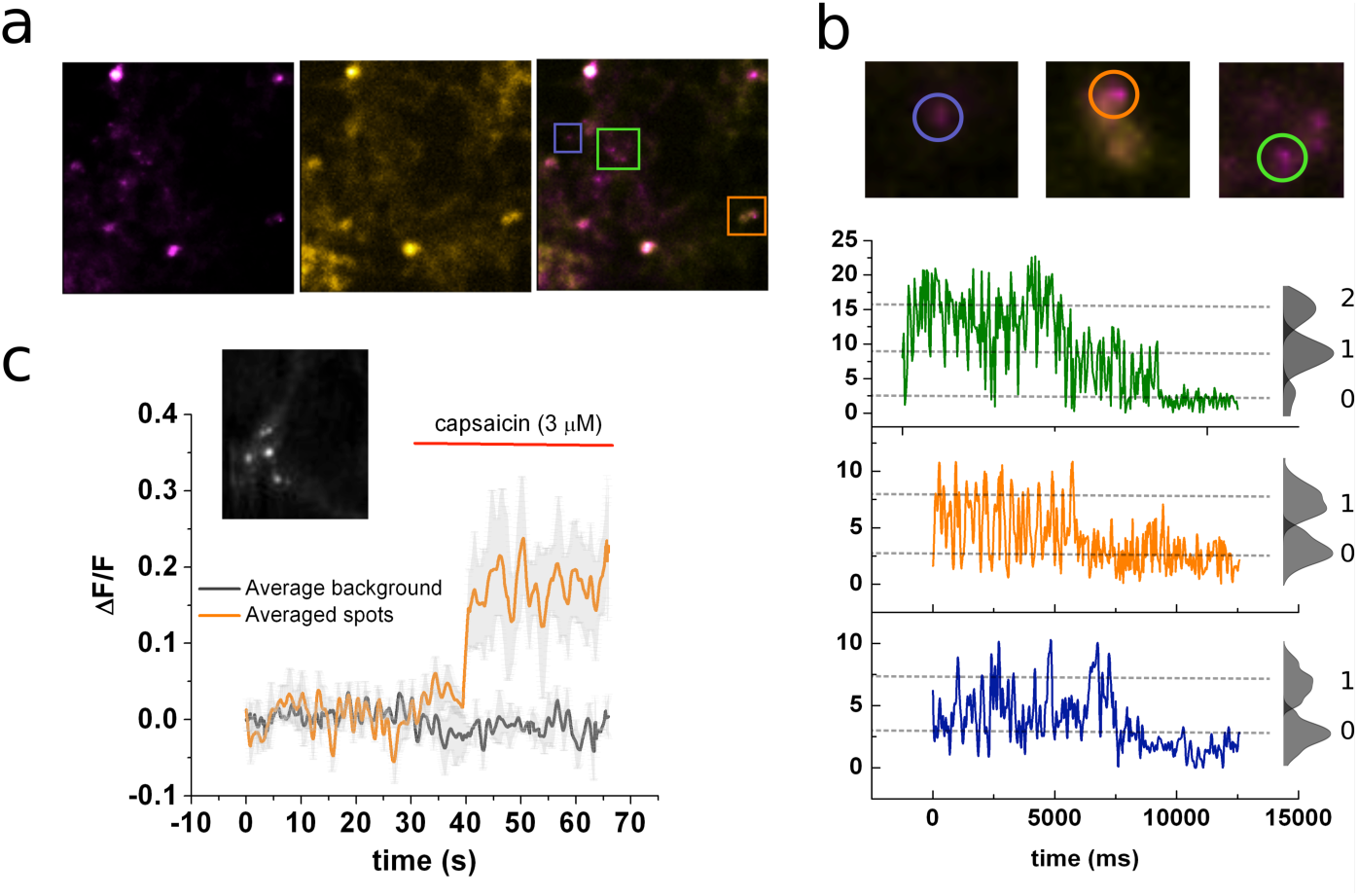
Identification of channels expressing a single coumarin emitter. **a.** Representative TIRF images of the 7:1 transfected cells. Images correspond to the average of 100 frames. Colocalization of coumarin diffraction-limited puncta with YFP signal is marked by colored squares. Bar correspond to 2 μm. **b.** Representative traces of photobleaching coumarin-positive puncta. Top images correspond to zoomed ROIs depicted in (**a**). Color-coded traces depict the time course of the fluorescence signal used later for the identification of individual puncta that bleaches on a single step. **c.** Ensemble average of the signal recorded from multiple spots from the cell’s membrane showed at the inset. Image corresponds to the average of 80 frames taken on the coumarin channel (ex405nm/em450nm).

### Activation kinetics obtained from the optical signal

Stationary noise analysis was next performed based on the autocorrelation (C_(Δt)_) of the fluorescence signal (figure 3a). The auto covariance function can be used to calculate the correlation between values of a noisy signal taken at increasing time intervals Δt. In this regard, C_(Δt)_ will be higher for small values of Δt (i.e. data points close to each other) and this function rapidly decays as Δt become larger. In the present example, this analysis allows for the identification of a characteristic time constant (τ) of the exponential decay describing the autocorrelation of the signal. The obtained τ has an inverse relationship to the rates defining the equilibrium of the reaction between open and close states (Anderson and Stevens, 1973; Zingsheim and Neher, 1974). The calculated autocorrelation function used discriminates well between the three conditions tested (i.e background, no capsaicin, capsaicin present in the media). The background signal is obtained from cells exposed to the coumarin amino acid but not the TRPV1^TAG^ cDNA. This condition generated noise with a single, rapid time constant (τ = 18 ms) that was easily separated from the slower time constants of the signal coming from the encoded coumarin conditions. Incubations with supersaturating capsaicin concentrations (3 μM) changed the slow time constant from 1360 ± 287 ms to 580 ± 182 ms (n=17; figure 3a), consistent with the possibility that the encoded coumarin is reporting on an agonist-induced increase in τ. Such a result can be interpreted in multiple ways: i) the stabilization of the open state after ligand binding (i.e. longer bursts of activity); ii) the destabilization of the close conformation (i.e. shorter close states); iii) or a combination of both, as demonstrated previously for TRPV1 channels (Hui et al., 2003).

In order to begin to distinguish between these possibilities, we examined unitary transitions in the optical data (figure 3b; see methods). To do so, we inferred that the signal minima (i.e. L_0_) would correspond to the magnitude of background signal and corroborate that fluorescence transitions drop from a given maxima (L_1_) to the observed minima (L_0_) as expected (figure 3b, inset). An analysis of the idealized signal shows that the agonist does not affect the amplitude of the signal, but rather it promotes a shortening of the mean time of the state we defined as L_0_, shortening from 398 ± 133 ms to 156 ± 52 ms (n=7; figure 3 c and d). This is in agreement with previous data showing that the main gating effect of capsaicin is the shortening the close state, instead of prolonging the duration of the open state (Hui et al., 2003), and suggests that coumarin at position Y671 directly describes the transition between open and close states of the channel. Given the physic-chemical properties of coumarin, the data suggest that position Y671 is more water exposed in the closed state, and thus the decrease in fluorescent output, compared to the open state (Wagner, 2009). By comparison, TRPV1 structural data place W426 in an invariant solvent exposed region, thus, the open-close transition should not induce evident changes in local solvation (Gao et al., 2016). To test this possibility directly, TRPV1 W426^Coum^ channels were optically examined (figure S1a). Electrophysiological analysis confirmed that W426^Coum^ is a functional channel (figure S1b). Further, consistent with the structural prediction, substitution at W426^coum^ lacked capsaicin dependent fluctuations (figure 3e). On the other hand, P_L1_ of position 671^coum^ reported fluorescent changes are well fit by a capsaicin dose response curve (figure 3e and figure S2). Lastly, when the fluorescence data from Y671^coum^ was normalized and compared with the electrophysiological data taken from whole cell macroscopic recordings, we observed a close match between the calculated Ec_50_ for optical and electrophysiological data, 0.33 μM versus 0.43 μM, respectively (figure 3f). Taken together, the data and companion biophysical analysis suggest a continuum of coordinated motions within the pore, with the Y671^coum^ optical data closely matching a transition to the open, conducting conformation in the selectivity filter.

**Figure 3.**
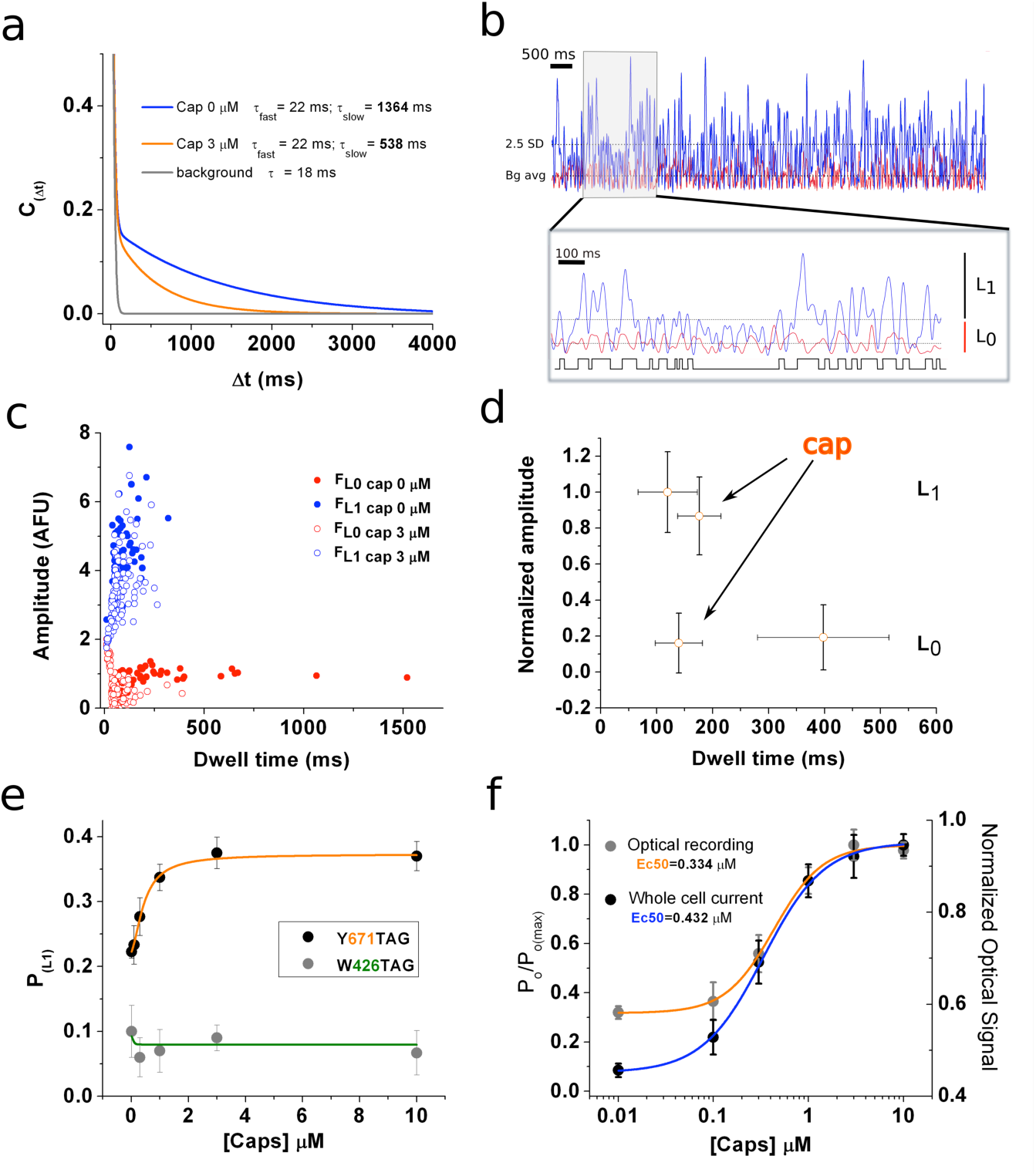
Optical studies of ion channel gating. **a.** Stationary noise of the single spots, each curve corresponds to the averaged fitted curves for the different conditions (n=13). Autocorrelation function was obtained for background (gray), capsaicin free (blue), and 3 μM capsaicin (red). A clear change in the slow time constant (τ_2_) can be observed. **b.** Analysis of the fluorescence fluctuations. Superimposed traces of signal (blue) and background (orange). The threshold for single channel opening is highlighted (2.5 SD, dotted line). The gray shaded box indicates the zoomed in portion of the recording that is presented on the inset on top. Same dotted lines indicating background average and threshold are shown. An idealization of the zoomed trace is shown in black. **c.** Amplitude versus dwell-time plot shows a shortening of mean time L_0_. **d.** Plot summarizing datasets of amplitude versus dwell time at 0 and 3 μM capsaicin (n=11). Error bars correspond to SEM. **e.** Dose-response curve showing changes in P_L1_, calculated from fluorescence fluctuations, for the two positions tested. **f.** Normalized dose-response curves for fluorescence fluctuations (orange) and whole cell electrophysiological response (purple) showing a good correlation of the Ec_50_ obtained under these different experimental procedures. The electrophysiological response was tested at 140 mV in the presence of the different concentrations of capsaicin. Error bars correspond to SEM.

### Side chain rearrangements within the pore

The observed changes in fluorescence can be explained by a change in solvation and/or in the coordination of the hydroxyl group of the dye (Wagner, 2009). To explore motion of Y671 at an atomistic level, we analyzed molecular dynamics (MD) trajectories of the TRPV1 closed and open states (about 750 ns each; figure 4a) obtained in our previous work (Kasimova et al., 2017). These two trajectories correspond to the equilibration of the capsaicin-bound structure (Cao et al., 2013), in which we either hydrated the cavities present between the S4- S5 linker and the S6 helix, or left them empty. We have previously shown that hydration of the peripheral cavities stabilizes the closed conformation of the channel, while initialization of the simulation with empty cavities triggers opening (Kasimova et al., 2017).

First, we compared the conformations that Y671 adopts in the closed and open states of TRPV1 (figures 4a and b). In the closed state, the representative conformation shows that three out of four tyrosine residues are oriented perpendicular to the membrane and do not contact each other (figure 4a). Instead, they create transient hydrogen bonds with F640 (backbone) of the selectivity filter. In the open state, the four-tyrosine residues reorient parallel to the membrane, forming a ring around the central pore (figure 4b). The altered orientation of these tyrosine residues in the two structural states affects hydration of the central pore (figure 4c). Surprisingly, the region of the pore surrounded by the four Y671 moieties is more hydrated in the closed than the open state (figure 4c). Furthermore, the hydration of Y671 is different in the two conformations: in the open channel, one face of the phenyl ring is exposed and the other one is partially buried by F640 and M644, while in the closed state both faces of the Y671 side chain are exposed to water. In order to more quantitatively describe Y671 hydration in the open and closed conformations the corresponding solvent accessible surface area (SASA) was calculated; see methods for details.

**Figure 4.**
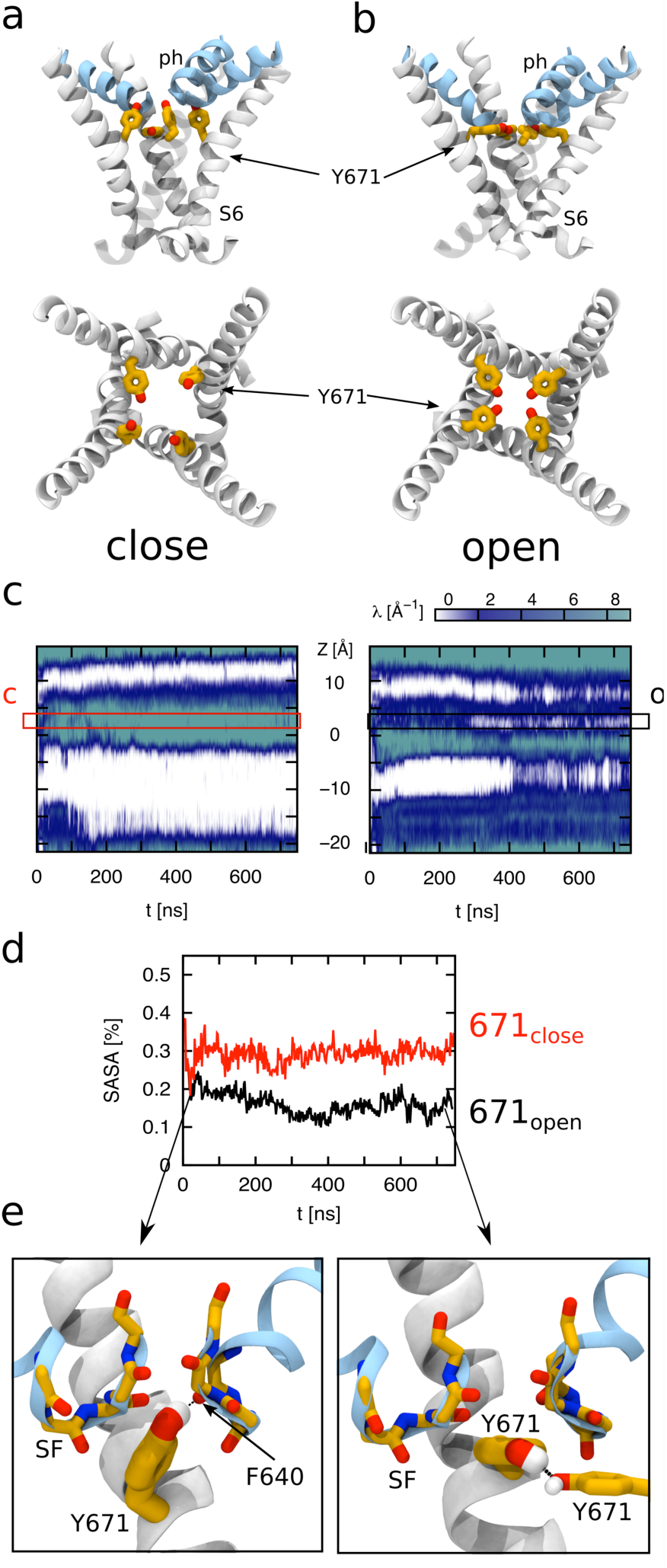
Molecular Dynamics simulations to calculate hydration of Y671. MD simulations of TRPV1 channels in the open (**a**) and closed (**b**) states. The upper and lower panels correspond to the side and top views on the S6 helices bundle (white), respectively. The pore helices (PH) are shown in cyan. The Y671 residues are represented as yellow sticks. Note that in the open state, the four Y671 residues create a ring oriented parallel to the membrane plane (perpendicular to the pore principal axis); while in the closed state, three of these residues do not interact with each other and are oriented perpendicular to the membrane plane (parallel to the pore principal axis). **c.** Hydration of the channel central pore. The left and right panels correspond to the close and open states, respectively. The y-axis of the plot denotes the z-axis of the system (parallel to the channel central pore). The origin of the axis is set at the center of the pore domain transmembrane part. The Y671 residues are located at ∼ 2.5 Å. Note that in the open state, at this level the water density is often interrupted starting from ∼ 280 ns, while in the closed state this density is continuous. **d.** The y-axis denotes the solvent accessible surface area (SASA) of Y671. The black and red curves correspond respectively to the open and closed states as indicated. Note that in the closed state the Y671 SASA is approximately two times larger compared to open starting from ∼ 300 ns. **e.** The panels show zoomed view on the Y671 residue in the close (left) and open (right) states. In the open state, Y671 establishes a hydrogen bond with Y671 of the adjacent subunit; while in the closed state, this residue interacts with the backbone of F640. The backbone of the selectivity filter (SF) is represented as sticks.

We found that the average Y671 SASA is larger in the closed state compared to open: in the closed state, about 30% of the Y671 surface area is exposed to water, while in the open state this value is approximately two times smaller (figure 4d). The evolution of the Y671 SASA along time reveals that the two MD trajectories start to diverge quite early: in particular, in both conformations, the Y671 SASA drops down from ∼ 35% to ∼ 20% at the beginning of the equilibration; then, after ∼ 30 ns, in the closed and open states respectively, the SASA either increases up to ∼ 30% or continues to decrease until it reaches an equilibrium value of ∼ 15% (figure 4d). Further, when local interactions are viewed in more detail, it can be observed that while the open state displays the side chains of Y671 residues engaged in mutual hydrogen bonding interactions, they establish vdW interactions with nearby residues F640 and M644, whose side chains are in between the selectivity filter (SF) backbone groups of two adjacent subunits (figure 4e). In the close state, F640 and M644 residues still interact with Y671 as in the open state; however, the different orientation of the side chain phenyl groups promotes a rigid displacement of the pore helix toward the extracellular side of the pore, narrowing the diameter of the selectivity filter (figure 4a and e). Overall, the MD simulations suggest that Y671 hydration decreases significantly as channels transition from the conducting to the non-conducting conformation, a notion that is in agreement with optical data from Y671-courmain channels.

## DISCUSSION

Previously, the incorporation of f-ncAAs and their use as reporters of membrane protein activity was achieved only in *Xenopus l*aevis oocytes, where macroscopic activity of Kv channels was recorded by the use of the f-ncAA ANAP (Kalstrup and Blunck, 2013). Additionally, a misacylated tRNA in the oocyte expression system have been used to encode BODIPY into nicotinic acetylcholine receptors, in order to examine stoichiometry of static protein at the plasma membrane with single molecule resolution (Pantoja et al., 2009). Recently, organic dyes from the cyano family of single molecule probes were encoded in a model chloride channel, ClC-0, (Leisle et al., 2016). Moreover, single molecule imaging of gating events have been previously reported for purified bacterial ion channels labeled with a cysteine-reactive dyes that could be observed when incorporated into an artificial bilayer system (Blunck et al., 2008) or liposomes (Wang et al., 2015). Thus, our imaging approach represents, to the best of our knowledge, the first direct optical recording of discrete gating-related events performed in a live cell.

Ion channel gating occurs in the range of hundreds of μs and it is one of the fastest protein-dependent processes within cells (Hille, 2001). This rapid time-frame imposes intrinsic technical barriers in the detection of a limited number of photons coming from few emitting molecules and the consequent low signal-to-noise ratio under these conditions (Ha, 2014). Given the design of our imaging setups and our experimental conditions (i.e. transient overexpression) we are bound to either the low (64 × 32 pixel) spatial resolution of the ns-time scale sampling of the SPAD imager (not allowing us to discriminate single emitters), or to a low signal-to-noise ratio restricting us to image at a maximum of 500 Hz (ms-time scale) for the individual diffraction-limited spots observed by an EM-CCD camera. As reported previously (Blunck et al., 2008), we observed that the latter approach is useful for conformational transitions on the order of 2-5 ms, thus barring optical access to the fast intra-burst channel activity which occurrs in tens of μs. Therefore, the unitary transition analysis presented here was limited to the description of each burst as a single opening at the expense of missing activity during short openings. None the less, such an analysis, while lacking detail on unitary transitions, is a valid approach to estimate mean open and closed channel durations. The ideal experimental system would allow for sparse location (∼1 micrometer) of the emitter-expressing receptors at the plasma membrane, so that the photon count profile of individual molecules can be obtained with the SPAD imager with nanosecond resolution. However, in the absence of such technical advances, standard biophysical approaches are in place to obtain insights on unitary gating events from an expressed population of channels. Interestingly, the similar rate constants obtained by noise analysis of the optical signal and from the subsequent modeling of the unitary transitions, suggest a strong internal consistency of data.

Another limitation to consider in the analysis is the intrinsic blinking of the dye. In the present case this blinking represents roughly 12% of the P_L1_ transitions and obfuscates efforts to obtain absolute values for the open probability. Our solution to bypass this issue was to linearly subtract this “background open probability” post-hoc and normalize the data to its maximal theoretical response. Still, even with this coarse approximation, the optical data is in surprising agreement with whole cell electrophysiological recording of channel gating. Further, this approach allowed for the direct measurement of the steady state activity of membrane receptors undergoing fast molecular rearrangements. Specifically, we observed that Y671^coum^ experienced a change in solvation as part of the process of pore rearrangement associated to channel’s opening. This was confirmed by our molecular dynamics calculations of solvent accessibility within the channel’s permeating pathway, showing a 15% decrease of Y671’s hydration in the open state conformation. Finally, in support of the overall analysis, the optical measurements closely overlaps with the dynamic range of burst activity observed in single channel recordings in response to capsaicin (i.e. 0.1 to 1 μM) (Hui et al., 2003).

Taken together, the observations presented here have shed new insights into the gating mechanisms of TRPV1 channels. Specifically, the data point to explicit conformational dynamics at the resolution of individual side-chains within the channel selectivity filter that report on the open to closed transition. It has been suggested previously that I679, a residue located near the inner bundle crossing, might act as a hydrophobic plug in the close state (Palovcak et al., 2015; Poblete et al., 2015), and that this barrier together with a second constriction, close to the selectivity filter, need to be released for the channel to fully open (Gao et al., 2016; Kasimova et al., 2017). This gives rise a possible gating scheme whereby the orientation of the TRP domain helix (TDh), covalently bound to the S6 helix and interconnected to the S4-S5 linker, will be affected by capsaicin by altering the interaction with the latter and transducing the motion trough the former where the pair N676-Y671 might play a role as modulators of the relative position of the selectivity filter (Gregorio-Teruel et al., 2015; Kasimova et al., 2017; Steinberg et al., 2014; Teng et al., 2015). In the present work we show no evidence of such a link but only the state-dependent changes at Y671. Therefore, the possibility still remains that the movement of Y671 is induced by capsaicin binding using an alternative pathway, which may not include the participation of the TDh, but rather the S5 helix.

## METHODS

### Cell culture

HEK293T cells, were cultured in DMEM (Gibco Inc.) supplied with 10% FBS (Gibco Inc.). Cells were prepared at (60-70)% confluence and transfected with lipofectamine 2000, by following the instructions from the provider (Life technologies). Often the target gene was transfected 2-3 hours after the initial transfection of the pair tRNAtag/CoumRS (Steinberg et al., 2016). The ncAA was added to cell cultures to a final concentration from 0.5 to 5 μM, together with the second transfection procedure. *<u>Imaging</u>*: To simplify the analysis of the optical data, we transfected a mixture of TRPV1^YFP^/TRPV1^TAG^ in a 7:1 ratio in an attempt to drive the system to incorporate as few coumarin molecules per tetramer as possible. 24 hours after transfection of the target gene, cells are disaggregated and plated on poly-L-lysine treated glass covers. The f-ncAA-containing media was removed at least 12 hours prior the experiment allowing cells to clear the soluble ncAA. Generally the cells are recorded 36-48 hours after the second transfection. *<u>Electrophysiology</u>*: Cells were disaggregated 2-3 hours before patching, and incubated with normal media.

### ***Synthesis* <u>of TyrCoum</u>**

L-(7-hydroxycoumarin-4-yl) ethylglycine was obtained as described before (Wang et al., 2006).

### Solutions

Ringer solution for imaging experiments contained: 140 mM NaCl, 8 mM KCl, 8mM HEPES, and 1 mM MgCl_2_ at pH 7.4. Electrophysiologycal solutions contained: *External*: 145 mM NaCl, 2 mM CaCl_2_, 5 mM KCl, 10 mM HEPES, 10 mM glucose, pH 7.4; *Internal*: 135 mM CsF, 5 mM KCl, 2 mM MgCl_2_, 1 mM CaCl_2_, 4 mM EGTA, 20 mM HEPES, pH 7.4.

### Molecular modeling

To analyze Y671 hydration at the atomic level, we used the molecular dynamics (MD) trajectories obtained in our previous work (Kasimova et al., 2017). These trajectories were generated starting from the cryo-EM structure of the TRPV1 capsaicin-bound state (Cao et al., 2013). In one simulation we initialized the system by inserting several water molecules (from 4 to 6) inside the channel peripheral cavities (located between the S4- S5 linker and the S6 C-terminus); in the second simulation we left these cavities empty. During the equilibration (about 750 ns), the molecular structures converged, respectively, to the closed and open states. To estimate water density along the central pore, we have implemented the following strategy. First, using HOLE software (Smart et al., 1996), we calculated the pore radius profile for each instantaneous configuration along the trajectory with a stride of 1 ns. Then, for the same set of frames, we calculated the three-dimensional histograms of water occupancy using the Volmap tool of VMD (Humphrey et al., 1996). Finally, we integrated the water occupancy in the XY plane (perpendicular to the channel central pore) using the pore radius profile as a boundary of the integration domain. The solvent accessible surface area (SASA) of Y671 in the closed and open states was estimated using the following procedure. From the MD trajectories we extracted a set of sub-trajectories, each containing 10 frames taken with a stride of 0.2 ns. For each sub-trajectory, we computed the three-dimensional histogram of atomic occupancy (for all the atoms except water) using the Volmap tool of VMD (Humphrey et al., 1996). We then used this map to define a molecular surface. To this end, we first discretized the map by assigning a value of 1 or 0, depending on whether or not the local occupancy is larger than a preset threshold. We considered all the bins with a value of 1 and located 1.5 Å away from a bin with a value of 0. Finally, the Y671 SASA was calculated as the overlap between the solvent accessible surface and the Y671 residue. Note that the SASA was normalized to the Y671 maximal surface area.

### Optical recordings

Cells were imaged using an inverted Olympus IX71 microscope main body and through-the-objective TIRF mode. Both 405 nm and 473 nm solid-state lasers (Coherent) were used to excite coumarin and YFP, respectively. Laser beams were focused to the backplane of a high-numerical aperture objective (Olympus 60X, N.A. 1.49, oil) by a combination of focusing lens. Fluorescence emission was collected by an Andor iXon^EM^ + 860 EM-CCD camera (Andor/Oxford Instruments, Ireland), after passing through an emission filter according to each acquisition wavelength band of interest (coumarin:450/70 nm; YFP:540/40 nm; Semrock, US). The light coming to the sample was controlled by a 12 mm mechanical shutter (Vincent associates, US), however, all measurements were performed under continuous light. All imaging experiments were done at room temperature (20-22°C). For *<u>localization and co-localization</u>*, images were recorded between 10-100 ms intervals (100-10 Hz). For *<u>autocorrelation and unitary fluctuation analysis</u>*, images were acquired at 2ms intervals (500 Hz). Laser and focus control was performed using micromanager. Acquisition and digitalization was done with Andor Solis software (Andor/Oxford Instruments, Ireland). *<u>Photon count:</u>* was made by using a single-photon avalanche diode (SPAD) photon-counting camera (64×32 pixels array), which is space correlated with the EM-CCD camera, thus allowing for the location of limiting diffraction puncta. A continuous light stimulation of 1 μs duration was used to excite coumarin emitters and photons were collected during that period. Laser TTL triggering and piezoelectric focus control (PIFOC-721, PI, Germany) was performed using Micro-Manager software (Vale Lab, UCSD, US). Acquisition and digitalization was done through a custom code written in C++ (SPADlab at POLIMI).

### Image analysis

For *<u>localization and co-localization</u>* the set of frames was averaged to increase signal-to-noise ratio. Noise analysis of the steady-state signal by autocorrelation was successfully employed in the past to investigate the kinetic properties of ion channel unitary events from recordings with low signal-to-noise ratios (Anderson and Stevens, 1973; Zingsheim and Neher, 1974). For *<u>autocorrelation</u>*, post acquisition Gaussian digital filter was used at 1/4^th^ of the acquisition frequency. The background signal was averaged and the standard deviation (SD) calculated. *<u>Unitary fluctuation analysis</u>*. To analyze the fluctuations of fluorescence we defined 2.5 SD of the background signal as a threshold to discriminate from the two regimes, basal level (L_0_) and higher signal level (L_1_). We reasoned that fluorescence fluctuations, intrinsic to the probe, might interfere with our calculations; therefore we measured soluble coumarin deposited on glass coverslip. From the latter we calculate a 12 % of positive transitions, a value we linearly subtract from all the individual points taken from the recorded mutants (Y671^coum^ or W426^coum^), a new corrected dataset having the apparent P_L1_ was obtained after substraction. Normalization of optical data was done by dividing signal amplitude by the maximal signal obtained (F/F_max_).

### HEK-293T electrophysiology

Whole cell currents were obtained from transiently transfected HEK-293T cells. Gigaseals were formed by using 2-4 MΩ borosilicate pipettes (Warner Instruments, Hamden, CT). Junction potentials (6-8 mV) were corrected for CsF/NaCl solutions. Seal resistance was 2-3 GΩ in all cases. Series resistance and cell capacitance were analogically compensated directly on the amplifier. The mean maximum voltage drop was about 4 mV. Whole cell voltage clamp was performed, and macroscopic currents from voltage steps were acquired at 20 kHz and filtered at 10 kHz, ramp protocols were acquired at 10 kHz and filtered at 5 kHz. Currents are presented in terms of densities. Data acquisition was made using an Axopatch 200B and a digidata 1320 (Molecular Devices Inc.), The acquisition and basic analysis of the data were performed with pClamp 8.0 software.

## Supplemental figures

**Figure S1.**
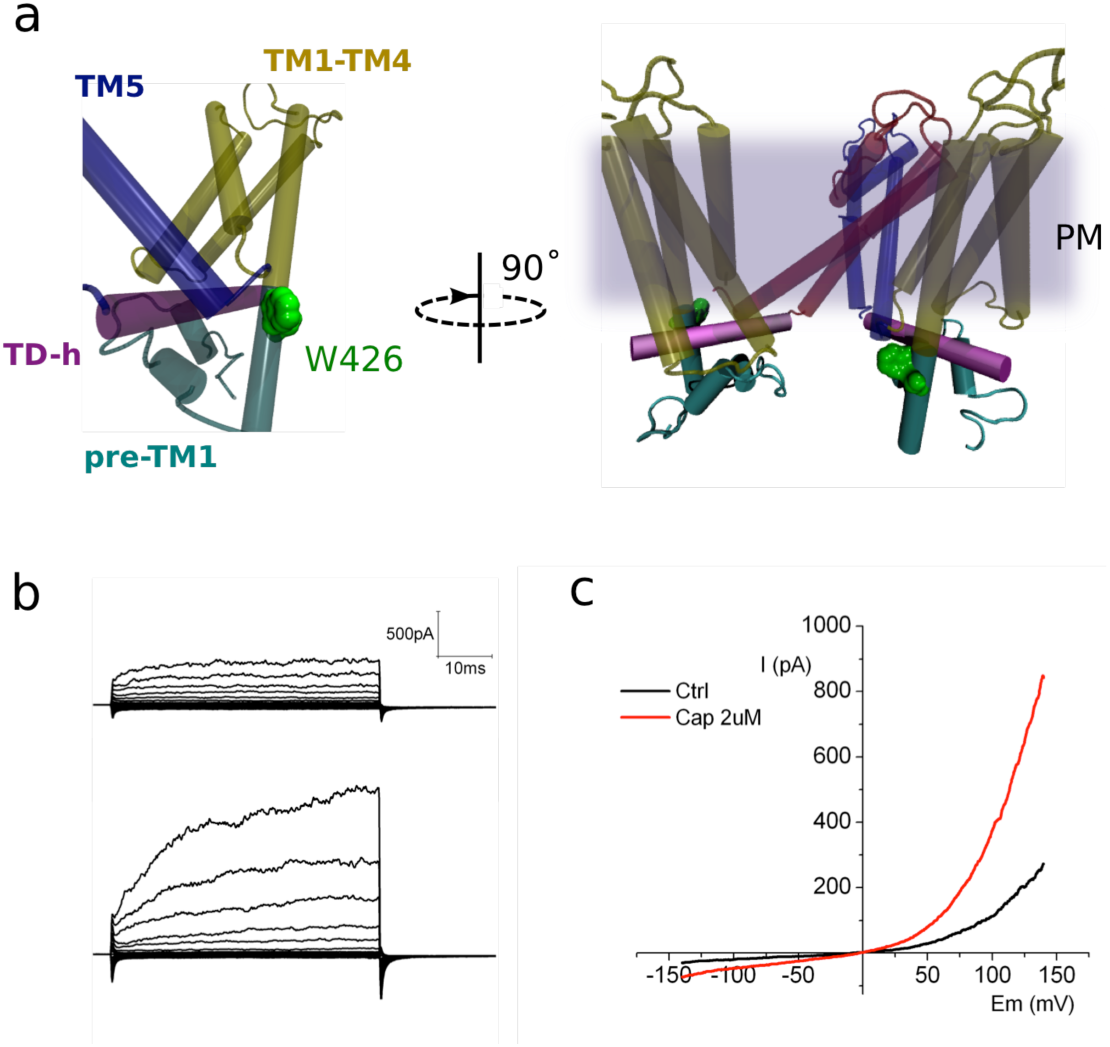
**a.** Two lateral views showing the relative position of W426, located on pre-TM1 region, outside the membrane. The different regions of the monomer are color coded as indicated. The image on the right depicts two subunits and the relative position of the channel with respect of the plasma membrane (diffuse gray). b. Representative traces obtained from transiently transfected HEK-293T cells subjected to voltage steps from -180 mV to +180 mV in 20 mV increments.

**Figure S2.**
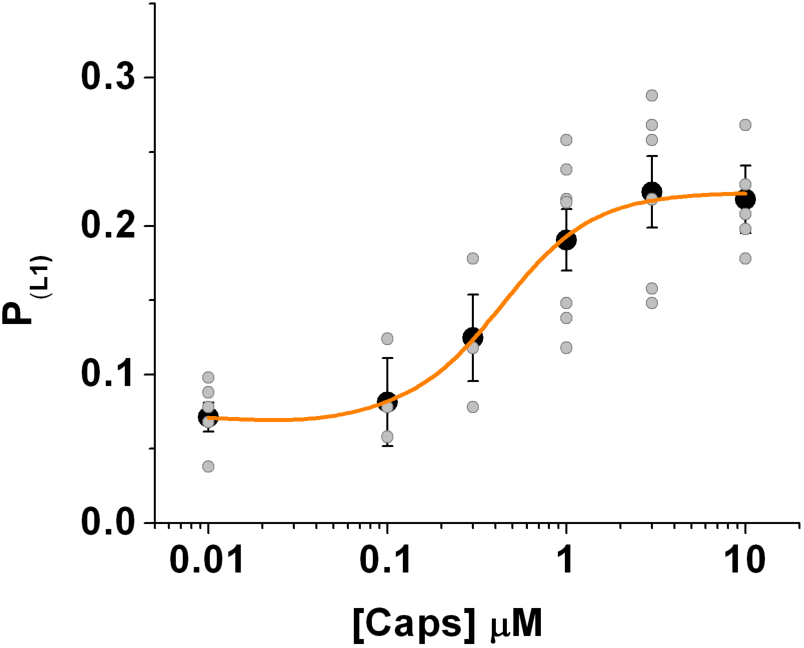
Distribution of the whole data set obtained from the fluctuation analysis of the fluorescence signal.

## Acknowledgments

X. Steinberg is a MECESUP and CONICYT fellow. This work was supported by FONDECYT grants 1110906 and 1151430 (SB), Anillo Científico ACT-1401 (SB), PCCI12023, DRI-CONICYT/CONACYT (SB and LDI). CAA is a supported by NIH/NIGMS (R01GM106569), is an American Heart Association Established Investigator (5EIA22180002) and a member of the Membrane Protein Structural Dynamics Consortium, which is funded by National Institutes of Health Grant GM087519 from NIGMS. LDI is supported by DGAPA-PAPIIT-UNAM grant IN209515. SB is part of CISNe-UACh and UACh Program for Cell Biology. VC acknowledges support from the National Institute of Health (R01GM093290) and National Science Foundation (ACI-1614804).

## Contributions

X.S., S.B, C.A.A and L.D.I. designed the project. X.S. performed experiments including biochemistry, molecular biology, and imaging. D.C-B. performed electrophysiological experiments. J.G. synthesize the ncAAs. M.A.K. and V.C. performed molecular dynamics simulations. M.A.K., X.S., E.L-G, and S.B. analyze data. F.V. developed the 64 × 32 single-photon counting camera. S.B., V.C., and C.A.A. provided reagents and materials. S.B, C.A.A. and L.D.I. wrote the paper.

